# Comparative transcriptomics identifies the transcription factors BRANCHED1 and TCP4, as well as the microRNA miR166 as candidate genes involved in the evolutionary transition from dehiscent to indehiscent fruits in *Lepidium* (Brassicaceae)

**DOI:** 10.1101/2021.10.29.466389

**Authors:** Lydia Gramzow, Katharina Klupsch, Noé Fernández Pozo, Martin Hölzer, Manja Marz, Stefan A. Rensing, Günter Theißen

## Abstract

**Background:** Fruits are the seed-bearing structures of flowering plants and are highly diverse in terms of morphology, texture and maturation. Dehiscent fruits split open upon maturation to discharge their seeds while indehiscent fruits are dispersed as a whole. Indehiscent fruits evolved from dehiscent fruits several times independently in the crucifer family (Brassicaceae). The fruits of *Lepidium appelianum*, for example, are indehiscent while the fruits of the closely related *L. campestre* are dehiscent. Here, we investigate the molecular and genetic mechanisms underlying the evolutionary transition from dehiscent to indehiscent fruits using these two Lepidium species as model system.

**Results:** We have sequenced the transcriptomes and small RNAs of floral buds, flowers and fruits of *L. appelianum* and *L. campestre* and analyzed differentially expressed genes (DEGs) and differently differentially expressed genes (DDEGs). DEGs are genes that show significantly different transcript levels in the same structures (buds, flowers and fruits) in different species, or in different structures in the same species. DDEGs are genes for which the change in expression level between two structures is significantly different in one species than in the other. Comparing the two species, the highest number of DEGs was found in flowers, followed by fruits and floral buds while the highest number of DDEGs was found in fruits versus flowers followed by flowers versus floral buds. Several gene ontology terms related to cell wall synthesis and degradation were overrepresented in different sets of DEGs highlighting the importance of these processes for fruit opening. Furthermore, the fruit valve identity genes *FRUITFUL* and *YABBY3* were among the DEGs identified. Finally, the microRNA miR166 as well as the TCP transcription factors *BRANCHED1* (*BRC1*) and *TCP FAMILY TRANSCRIPTION FACTOR 4* (*TCP4*) were found to be DDEGs.

**Conclusions:** Our study reveals differences in gene expression between dehiscent and indehiscent fruits and uncovers miR166, *BRC1* and *TCP4* as possible causes for the evolutionary transition from dehiscent to indehiscent fruits in *Lepidium*.

## Background

Flowering plants (angiosperms) form fruits to protect and disperse their seeds. Fruits come in many different types with different morphologies and different properties such as dry or fleshy, and dehiscent or indehiscent (Lorts and Briggeman 2008). There is a tremendous variation in fruit types both across and within different plant lineages (Dardick and Callahan 2014). However, the evolutionary mechanisms that enabled such dramatic shifts to occur, often in a relatively short period of time, remain largely unknown.

The crucifer family (Brassicaceae) includes a number of economically important plants such as cabbage, broccoli, mustard, radish, and turnips. The model plant *Arabidopsis thaliana* is also a member of this family (Initiative 2000). Typical fruits of Brassicaceae species are dehiscent, i.e. that the fruits open upon maturation to release the seeds. Dehiscent fruits also likely represents the ancestral fruit type of Brassicaceae (Hall et al. 2002). However, indehiscent fruits, i.e. fruits that only release the seed upon decomposition of the fruit, are found in many tribes distributed across the Brassicaceae phylogeny (Appel et al. 2003). The scattered distribution of indehiscent fruits indicates that this property evolved independently several times. This situation is mirrored in the genus *Lepidium* belonging to Brassicaceae: Species of this genus typically produce two-seeded dehiscent fruits, but the genus also includes species with indehiscent fruits (Al-Shehbaz and Mummenhoff 2011).

Brassicaceae fruits are composed of two fruit valves that are connected to the replum and enclose the developing seeds. Dehiscent fruits, such as those of *A. thaliana* and *Lepidium campestre* (also known as field pepperwort or field cress), form a well-defined dehiscence zone (DZ) at the valve margin (Mühlhausen et al. 2013). The DZ consists of the lignified layer, a stripe of lignified cells, and a separation layer, a region of small thin-walled cells (Spence et al. 1996; Rajani and Sundaresan 2001). During fruit ripening, the whole fruit dries and shrinks. Only the lignified structures stay rigid. Thereby a spring-like tension is created within the fruit. At the same time, the middle lamellae of the separation layer cells degenerate to form a pre-determined breaking zone at which the pressure tears the valves apart from the replum. Consequently, the fruit bursts open to release the seeds (Meakin and Roberts 1990, 1991; Spence et al. 1996). In contrast, the indehiscent fruits of the closely related *Lepidium appelianum* do not form a DZ. Instead, a continuous ring of lignified cells surrounds the seeds such that the fruit cannot open (Mühlhausen et al. 2013).

Much of the gene regulatory network underlying the proper formation of the fruit valves, replum and DZ has been elucidated in *A. thaliana* (reviewed in (Ballester and Ferrándiz 2017)). Establishment of the DZ requires expression of the two redundant MADS box genes, *SHATTERPROOF1* (*SHP1*) and *SHATTERPROOF2* (*SHP2*). The SHP1 and SHP2 proteins act as transcription factors and activate the basic helix-loop-helix protein-encoding genes *INDEHISCENT* (*IND*), *ALCATRAZ* (*ALC*) and *SPATULA* (*SPT*), and also autonomously contribute to DZ development (Liljegren et al. 2000; Liljegren et al. 2004; Rajani and Sundaresan 2001; Groszmann et al. 2011).

For correct fruit patterning, it is crucial that the expression of the *SHP* genes is restricted to the DZ. Three transcription factors contribute to this process: The MADS box gene *FRUITFULL* (*FUL*) which is expressed in the fruit valves (Gu et al. 1998; Ferrandiz et al. 2000), the BEL1-like homeobox gene *REPLUMLESS* (*RPL*) (Roeder et al. 2003), also known as *PENNYWISE* (Smith and Hake 2003), *BELLRINGER* (Byrne et al. 2003), *VAAMANA* (Bhatt et al. 2004), and *BLH9* (Cole et al. 2006) which is expressed in the replum, and the floral homeotic gene *APETALA2* (*AP2*) which also negatively regulates *RPL* (Ripoll et al. 2011).

Transcription factors controlling the expression of these negative regulators have also been determined. High levels of the C2H2 zinc finger proteins JAGGED (JAG) and the two closely related YABBY1 group proteins FILAMENTOUS FLOWER (FIL) and YABBY3 (YAB3) activate the expression of *FUL* (Dinneny et al. 2005). In contrast, lower levels of JAG/FIL/YAB3 expression promote expression of *SHP* genes. The expression of RPL is activated by the knotted1-like homeobox protein BREVIPEDICELLUS (BP) (Alonso-Cantabrana et al. 2007) whose gene is in turn activated by the C2H2 zinc finger protein NO TRANSMITTING TRACT (NTT) (Marsch-Martínez et al. 2014). *AP2* is negatively regulated be the microRNA miR172 (Ripoll et al. 2015).

Additionally, other factors which influence the size and the position of the DZ have been identified. The *WUSCHEL-RELATED HOMEOBOX gene 13* (*WOX13*) controls replum width and negatively regulates *JAG*/*FIL*/*YAB3* (Romera-Branchat et al. 2013). The auxin-response factors ARF6 and ARF8, which are regulated by miR167 (Zheng et al. 2019), activate miR172 together with FUL (Ripoll et al. 2015). The MYB protein ASYMMETRIC LEAVES 1 (AS1), likely in collaboration with the leucine zipper protein ASYMMETRIC LEAVES 2 (AS2), negatively regulates BP (Alonso-Cantabrana et al. 2007).

In general, proteins encoded by genes expressed in the replum often negatively regulate genes expressed in the valves and *vice versa*. Apart from the already mentioned interactions, this includes negative regulation of the replum gene *BP* by the valve proteins encoded by *JAG*/*FIL*/*YAB3*, and negative regulation of *JAG*/*FIL*/*YAB3* by the replum protein RPL (González-Reig et al. 2012).

In a previous study, we have shown that orthologues of the valve margin genes are expressed in a similar way in *L. campestre* (dehiscent fruits) as in *A. thaliana* fruits but that expression of the respective orthologues is abolished in the corresponding tissues of indehiscent *Lepidium appelianum* fruits (Mühlhausen et al. 2013). As parallel mutations in different genes are unlikely, we concluded that the changes in gene expression patterns are probably caused by changes in upstream regulators such as FUL, RPL or AP2.

To conduct a more unbiased approach to identify the genetic changes that lead from dehiscent to indehiscent fruits than the analysis of candidate genes, we have sequenced the transcriptomes of floral buds, flowers and fruits of both, *L. campestre* and *L. appelianum* in the present study. We have identified differentially expressed genes (DEGs) and differently differentially regulated genes (DDEGs) where the latter refers to genes for which the change in expression level between two structures is significantly different in one species than in the other. More DEGs were identified in flowers than in fruits and floral buds and a higher number of DDEGs was found in fruits versus flowers than in flowers versus floral buds. Cell wall synthesis and degradation are important processes for fruit opening as revealed by gene ontology (GO) analysis. The fruit valve identity genes *FRUITFUL* and *YABBY3* were identified as DEGs such that the possible cause for the evolutionary transition from dehiscent to indehiscent fruits in *Lepidium* may even be an upstream factor of these genes. Possible candidates are *BRANCHED1* (*BRC1*), an ortholog of which may determine whether dehiscent or indehiscent fruit develop on the dimorphic plant *Aethionema arabicum*, and *TCP FAMILY TRANSCRIPTION FACTOR 4* (*TCP4*) which may regulate *YABBY3*. These two genes were found to be DDEGs. Our study elucidates differences in gene expression patterns between dehiscent and indehiscent fruits and reveals *BRC1* and *TCP4* as possible causes for the evolutionary transition from dehiscent to indehiscent fruits in *Lepidium*.

## Results

### Overview of the RNA-seq analysis and transcriptome assembly

Sequencing resulted in an average number of reads per library of 56 Mio. for the mRNA and 12 Mio. for the small RNA (Table 1). An initial analysis of the data revealed contamination with sequences from thrips, likely due to infestation of the plants by these animals. Hence, we removed reads matching to the genome of the thrips *Frankliniella occidentalis* (González et al. 2018) as well as uncorrectable and unpaired reads and reads corresponding to organelle sequences. After this filtering step, 42 Mio. were retained for further analyses for the mRNA sample. For the small RNA sample many reads seem to be derived from organelle RNA. Hence, after removing uncorrectable reads and those matching to the *Frankliniella occidentalis* genome and organelle sequences, only 1.5 Mio. reads remained on average for the small RNA sample (Table 1).

**Table 1:** Number of reads obtained after sequencing and after correction and pruning steps.

| Experiment | Species | Structure | Replicate | Raw reads | Uncorrectable, unpaired reads removed | Thrips and organelle sequences removed |
| --- | --- | --- | --- | --- | --- | --- |
| mRNA | *L. campestre* | Bud | 1 | 56,364,306 | 47,437,984 | 42,661,682 |
|  |  |  | 2 | 52,626,578 | 44,103,956 | 42,345,608 |
|  |  |  | 3 | 46,984,896 | 38,458,226 | 35,556,348 |
|  |  | Flower | 1 | 53,973,840 | 45,071,412 | 42,388,254 |
|  |  |  | 2 | 54,176,184 | 43,661,258 | 40,639,670 |
|  |  |  | 3 | 47,473,062 | 37,866,352 | 33,766,726 |
|  |  | Fruit | 1 | 66,540,836 | 59,074,352 | 48,284,508 |
|  |  |  | 2 | 67,087,830 | 57,231,738 | 45,811,592 |
|  |  |  | 3 | 57,259,526 | 48,363,464 | 41,299,370 |
|  | *L. appelianum* | Bud | 1 | 61,044,624 | 51,099,508 | 47,872,858 |
|  |  |  | 2 | 57,117,360 | 45,964,748 | 41,954,354 |
|  |  |  | 3 | 56,803,334 | 47,261,624 | 42,970,772 |
|  |  | Flower | 1 | 54,109,008 | 45,806,166 | 41,765,762 |
|  |  |  | 2 | 60,218,542 | 49,851,062 | 46,492,188 |
|  |  |  | 3 | 59,056,126 | 49,511,932 | 46,039,554 |
|  |  | Fruit | 1 | 51,115,752 | 42,013,148 | 38,781,852 |
|  |  |  | 2 | 53,010,164 | 42,607,090 | 40,845,816 |
|  |  |  | 3 | 58,202,346 | 48,358,214 | 43,294,808 |
| smallRNA | L. campestre | Bud | 1 | 12,317,448 | 11,640,550 | 1,065,134 |
|  |  |  | 2 | 12,888,040 | 11,434,250 | 1,234,296 |
|  |  |  | 3 | 13,084,802 | 18,556,186 | 4,545,624 |
|  |  | Flower | 1 | 13,199,751 | 12,576,208 | 1,321,440 |
|  |  |  | 2 | 14,250,796 | 21,800,894 | 4,334,830 |
|  |  |  | 3 | 11,931,106 | 10,446,892 | 991,708 |
|  |  | Fruit | 1 | 12,600,710 | 10,038,752 | 908,384 |
|  |  |  | 2 | 12,171,462 | 10,194,186 | 430,032 |
|  |  |  | 3 | 12,245,803 | 15,075,818 | 1,955,400 |
|  | L. appelianum | Bud | 1 | 11,107,988 | 7,005,858 | 444,744 |
|  |  |  | 2 | 12,258,200 | 7,404,808 | 1,044,044 |
|  |  |  | 3 | 12,371,624 | 11,155,240 | 1,227,638 |
|  |  | Flower | 1 | 11,665,136 | 7,039,858 | 359,098 |
|  |  |  | 2 | 10,806,199 | 6,161,922 | 279,758 |
|  |  |  | 3 | 10,702,759 | 5,090,392 | 250,238 |
|  |  | Fruit | 1 | 11,180,533 | 10,409,022 | 1,667,408 |
|  |  |  | 2 | 11,088,927 | 10,542,716 | 1,354,590 |
|  |  |  | 3 | 12,727,467 | 16,904,122 | 3,445,396 |

Assembly using Trinity (Grabherr et al. 2011a) resulted in a total of 56,413 transcripts for *L. campestre* and 70,380 transcripts for *L. appelianum* after removing putative contaminant sequences but including potential splice variants or fragmentary sequences. The assemblies also contained chimeric sequences composed of two different transcripts which were likely a result of mis-assembly (Yang and Smith 2013b). Separation of chimeric sequences increased the number of transcripts to a total of 57,209 for *L. campestre* and 71,332 for *L. appelianum*. We used the Benchmarking Universal Single-Copy Orthologs (BUSCO) tool (Simão et al. 2015) with the dataset eudicotyledons_odb10 as reference to assess completeness of our transcriptomes. The BUSCO analyses revealed that 94.6% of the expected eudicotyledonous “near-universal single-copy orthologs” are present in our assembly of the *L. campestre* transcriptome while 94.3% of these BUSCOs are present in our *L. appelianum* transcriptome (Figure 1). It is common that some genes are fragmented in *de novo* assemblies. Hence, we analyzed the length distribution of our assemblies. For both species there are two peaks (Figure 2). One peak appears at a length of about 240 nucleotides and probably represents fragments. The other peak was found at a length of about 1,450 nucleotides which indicates that there are also a number of full-length transcripts.

**Figure 1:**
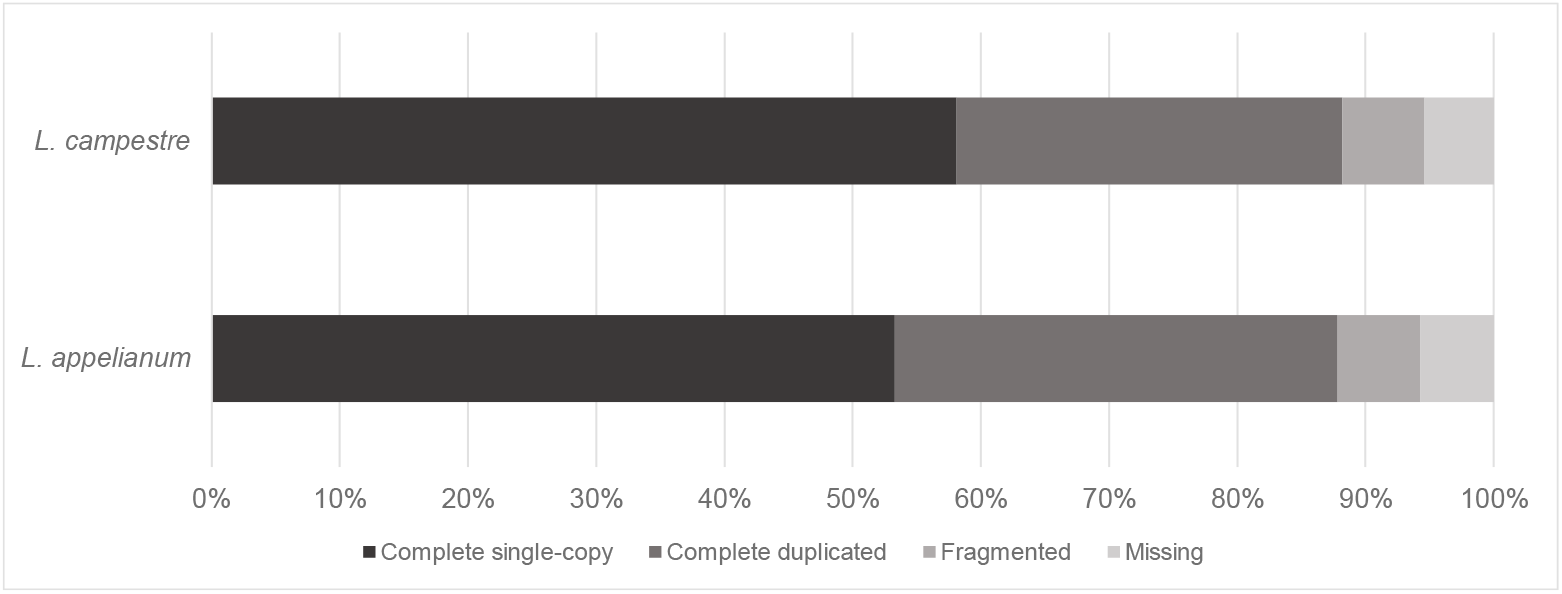
BUSCO completeness analysis. Transcripts from the *L. campestre* and *L. appelianum* assemblies were compared to 2121 Eudicotyledons reference orthologs for completeness assessment.

**Figure 2:**
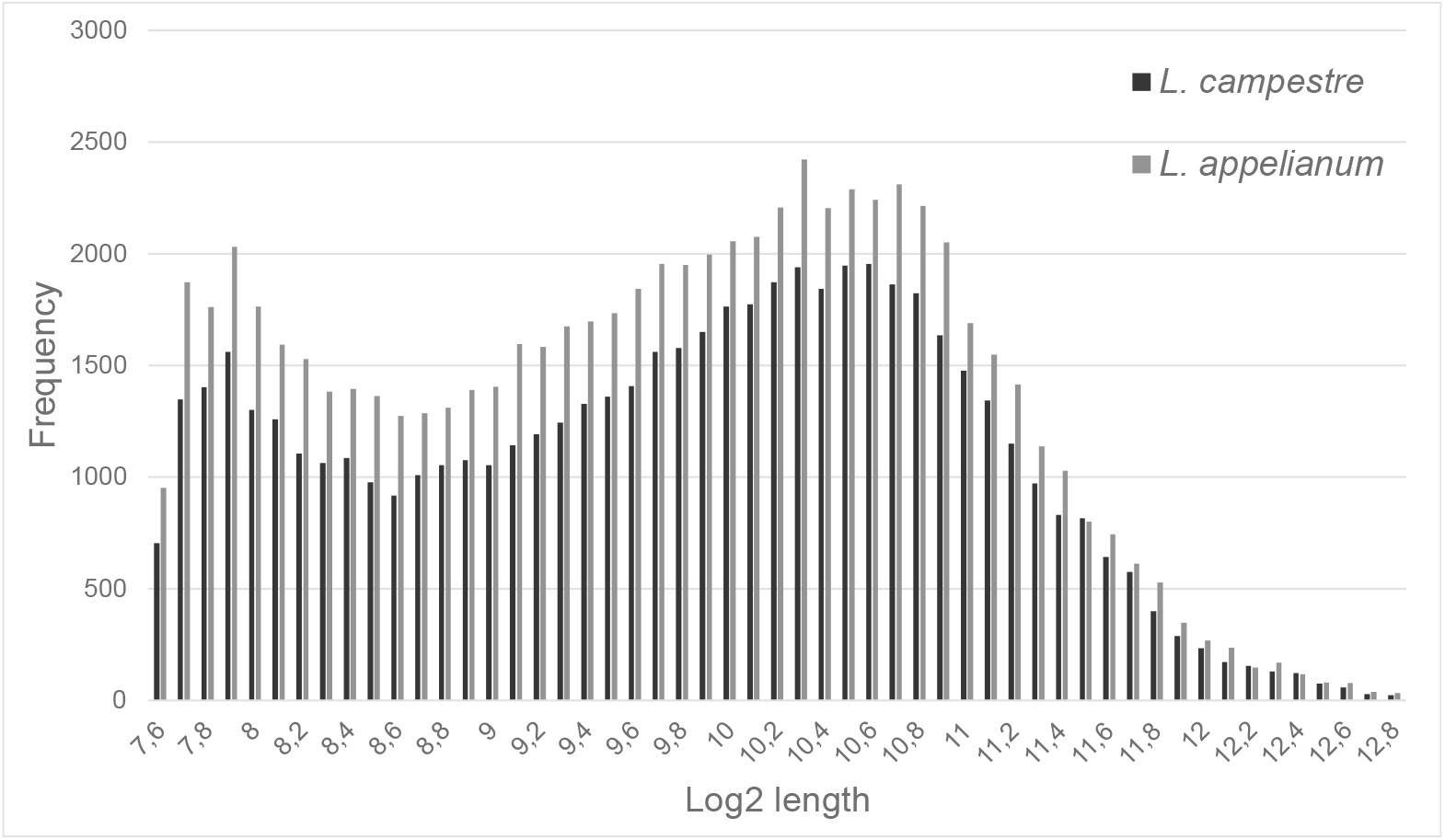
Transcript length distribution of the assembled transcripts of *L. campestre* and *L. appelianum*.

To detect conserved miRNAs, we mapped the small RNA reads onto the mature miRNAs of *A. thaliana* as provided by miRBase (Kozomara et al. 2019a). We found reads for 64 mature miRNAs belonging to 32 miRNA families in the *L. campestre* small RNA data (Table 2). Using ShortStack (Axtell 2013) and the *L. campestre* genome as available from NCBI, we identified three novel miRNAs. However, no putative target genes could be identified in the transcriptome of *L. campestre* using targetfinder (https://github.com/carringtonlab/TargetFinder). Our *L. appelianum* small RNA data contained reads of 60 mature miRNAs belonging to 30 miRNA families (Table 2). No novel miRNAs could be identified for *L. appelianum* using ShortStack and our transcriptome as reference “genome”.

**Table 2:**
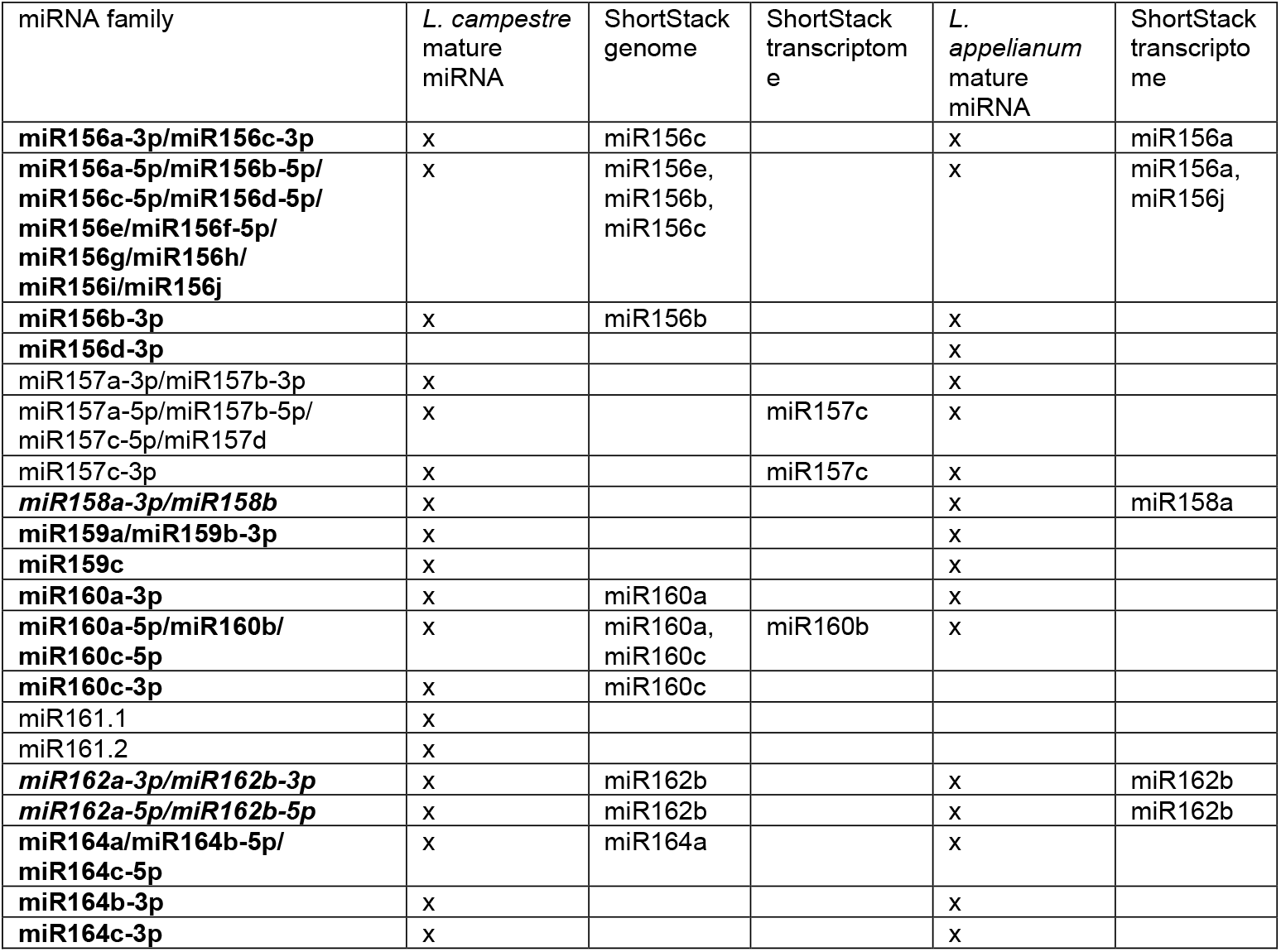
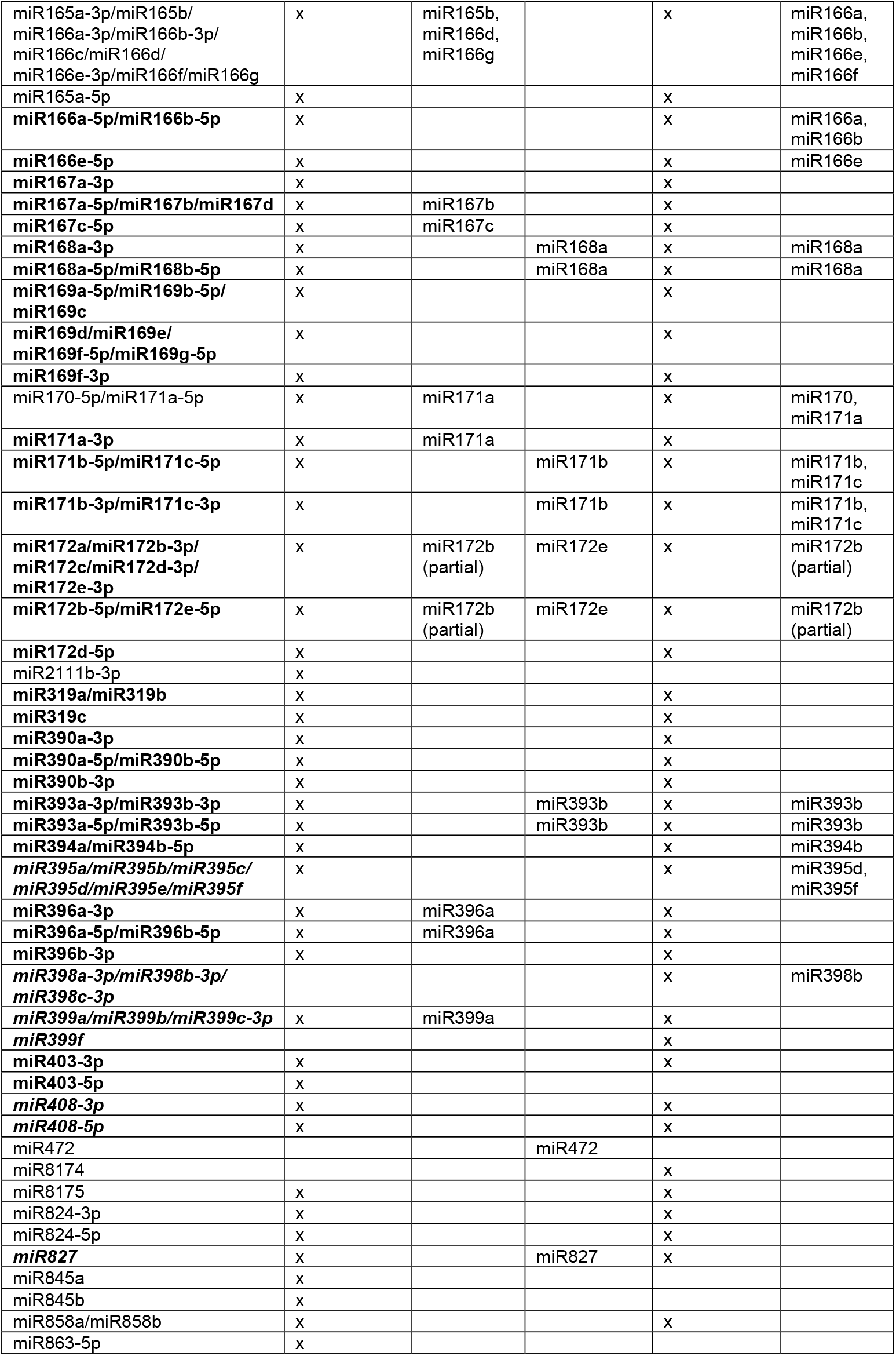
Mature miRNAs identified in short read data of *L. campestre* and *L. appelianum*. bold conserved miRNAs according to Chavez-Montez et al., 2014, bold and italic moderately conserved miRNAs according to Chavez-Montez et al., 2014

To assess completeness of our small RNA data, we compared our results to the set of conserved and moderately conserved miRNA families as identified by miRNA sample sequencing of vascular plants (Montes et al. 2014). For both species, we identified reads for all 16 miRNA families that were found to have originated before the emergence of eudicots and to be conserved across virtually all corresponding species. Furthermore, we found reads for 6 miRNA families in our *L. campestre* and 7 miRNA families in our *L. appelianum* small RNA data out of 21 miRNA families which were classified as conserved, although missing in a few corresponding species.

### Differential gene expression analysis

To conduct differential expression and regulation analysis, we identified putative ortholog pairs between the transcripts of the two *Lepidium* species as described in the methods section. We attained two transcriptome datasets, one for *L. campestre* and one for *L. appelianum*, each containing 17,755 transcripts and where each transcript in one species has exactly one putative orthologous transcript in the other species. We will refer to these transcriptome datasets as our ortholog-transcriptomes in the following. We reassessed completeness of our ortholog-transcriptomes and found that 89.2% of the BUSCOs remained in our ortholog-assembly for *L. campestre* while this value was slightly lower at 89.1% for our *L. appelianum* ortholog-transcriptome.

Reads were mapped independently to the corresponding ortholog-transcriptome and counted using HTSeq-count (Anders et al. 2015b). A principal component analysis was conducted based on the normalized number of reads mapping to the ortholog-transcriptomes. As expected, the replicates from the same species and structure clustered together (Figure 3). The species are separable based on first component which explains 54% of the variance while the structures are separable based on second component which explains 30% variance (Figure 3).

**Figure 3:**
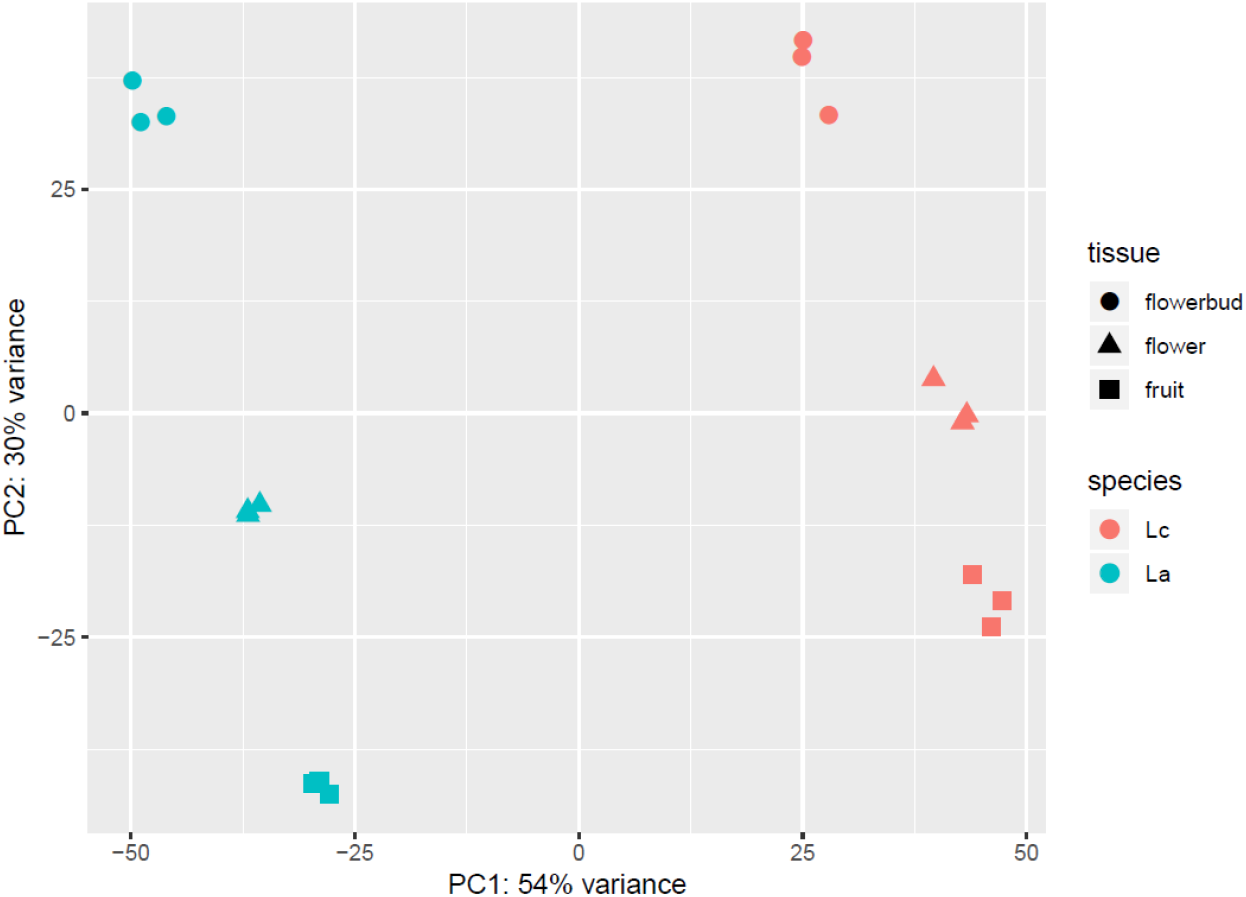
Principal component analysis of gene expression profiles of all samples. Samples from *L. campestre* are shown in red, samples from *L. appelianum* are shown in blue. Samples from floral buds are depicted by circles, samples from flowers by triangles and samples from fruits by squares. PCA shows separation of the two species and the different structures.

To learn more about the differences in fruit development between the *L. campestre* and *L. appelianum*, we analyzed expression in our ortholog-transcriptomes using the programs DESeq2 (Love et al. 2014a) and edgeR (Robinson et al. 2010a). We used a multi-factor design to not only be able to identify differentially expressed genes (DEGs) between the species in the same structure and between structures in the same species, but also to identify genes where the change in expression between the structures is different between the two species. We will refer to the genes identified in the latter analyses as differently differentially expressed genes (DDEGs).

DESeq2 generally identified more DEGs and DDEGs than edgeR, but there is a great overlap of genes identified by both programs (Figure 4). Only this overlap between the two methods will be considered in the following. More DEGs were observed between the same structure of the different species as compared to different structures of the same species. In *L. campestre*, there are similar numbers of DEGs between flower and bud as compared to fruit and flower. In *L. appelianum*, there are more than twice as many DEGs in flowers versus buds as compared to fruits versus flowers (Figure 4). When looking at DEGs in the same structure of the different species, the highest number of DEGs in observed in flowers, followed by fruits and buds.

**Figure 4:**
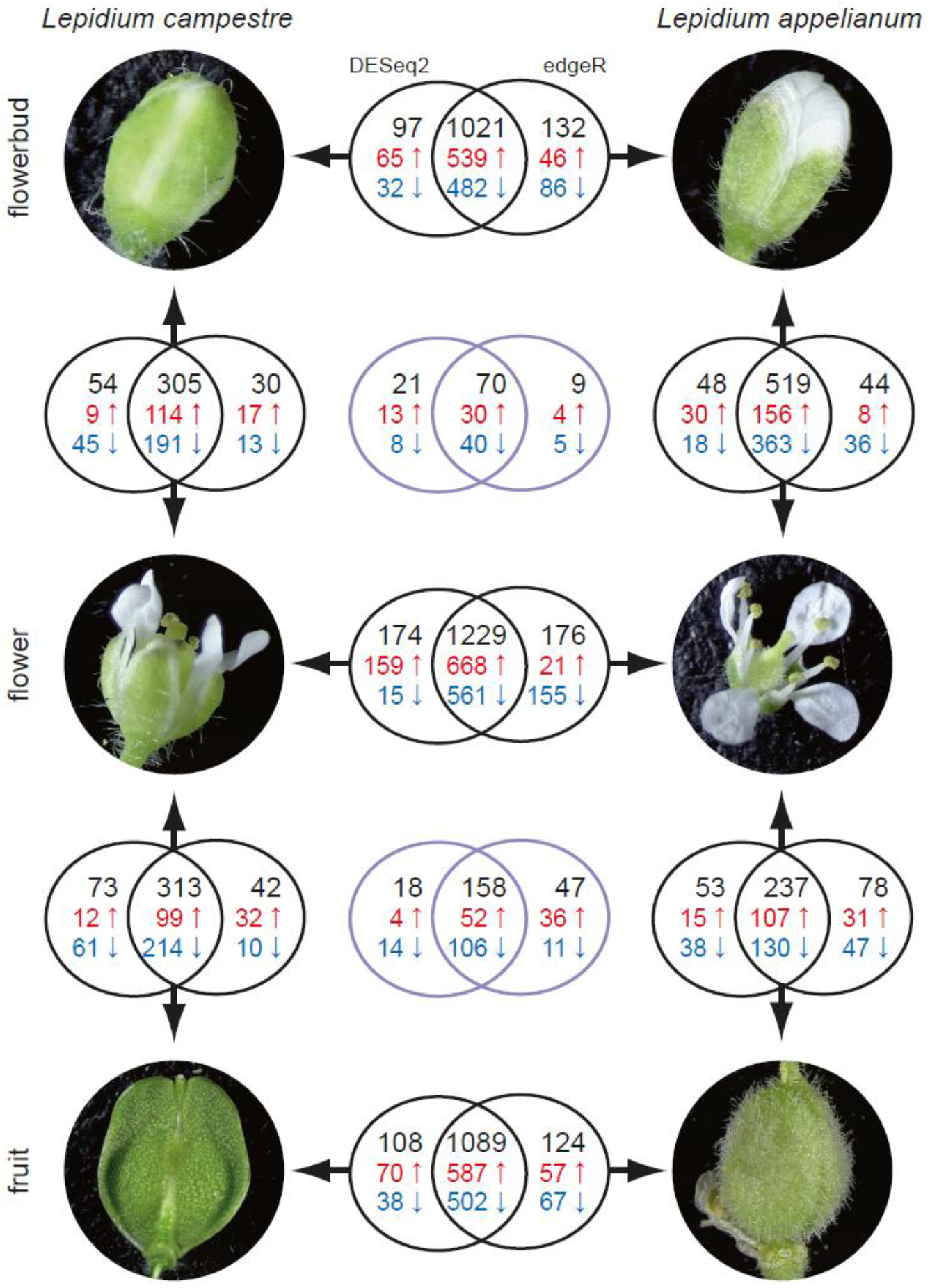
Venn diagrams of the DEGs and DDEGs between different species and different structures. The DEGs and DDEGs were called by the two programs edgeR and DESeq2. Venn diagrams between floral buds, flowers and fruits, respectively, of *L. campestre* and *L. appelianum* represent differentially expressed genes (DEGs) between the two species in the corresponding structure. Venn diagrams between different structures of the same species represent DEGs between those structures in the corresponding species. The two lavender Venn diagramms indicate differently differentially expressed genes (DDEGs) between flower and floral buds and between fruits and flowers, respectively, when comparing the two species. Black numbers in the Venn diagram correspond to all DEGs or DDEGs while red numbers represent up- and blue numbers represent downregulated genes.

We also analyzed DDEGs in our dataset, i.e. genes which had a significantly different change in expression in flowers versus buds and in fruits versus flowers, respectively, in *L. appelianum* as compared to *L. campestre*. These genes may have a significantly stronger up- or downregulation in *L. appelianum* as compared to *L. campestre* or these genes may be downregulated in one species and upregulated in the other species. We found 70 DDEGs in flowers versus buds and 158 DDEGs in fruits versus flowers when comparing the two species (Figure 4).

We applied the same methods for the identification of DEGs and DDEGs encoding miRNAs. First, we determined orthologs between the miRNAs based on the *A. thaliana* miRNAs they mapped to. For 56 mature miRNAs belonging to 28 miRNA families reads were found in the small RNA data for both species and these mature miRNAs could thus be used for differential expression analyses (Table 2). We will refer to this dataset as our ortholog-miRNAs. All 16 miRNA families that are conserved across virtually all species according to (Montes et al. 2014) and 6 out of 21 miRNA families which were classified as moderately conserved belong to our ortholog-miRNAs dataset. Mapping of small RNA reads, counting and differential expression analyses were done as described for the differential expression analysis of the ortholog-transcriptomes.

Only one miRNA was found to be encoded by a DEG or DDEG by both programs DESeq2 and edgeR. The miRNA homologous to miR165a-3p, miR165b, miR166a-3p, miR166b-3p, miR166c, miR166d, miR166e-3p, miR166f and miR166g of *Arabidopsis thaliana* (Reinhart et al. 2002) (they all only differ by one nucleotide), which we will refer to as miR165a-3p, was found to be encoded by a DDEG when comparing fruits and flowers. Targets of miR165 and miR166 are HD-Zip transcription factors like PHABULOSA, REVOLUTA and PHAVOLUTA (Rhoades et al. 2002).

### Gene Ontology and transcription factor analyses

A number of gene ontology (GO) terms (Ashburner et al. 2000; Gene Ontology Consortium 2021) of the category molecular function are significantly over- or underrepresented in the DEGs and DDEGs (Table 3). Among them, the terms protein binding (GO:0005515) and RNA binding (GO:0003723) were underrepresented in two datasets of DEGs. Interestingly, several GO terms related to cell wall synthesis and degradation, i.e. pectinesterase activity (GO:0030599), cellulose synthase (UDP-forming) activity (GO:0016760), polygalacturonase activity (GO:0004650) and hydrolase activity, hydrolyzing O-glycosyl compounds (GO:0004553) were overrepresented in different sets of DEGs.

**Table 3:** Gene ontology (GO) terms significantly over- or underrepresented in DEGs and DDEGs. Terms of the category molecular function of GO were analysed.

| Dataset | GO term | Fold enrichment | FDR |
| --- | --- | --- | --- |
| La vs. Lc bud | protein binding (GO:0005515) | 0.74 | 3.43E-02 |
|  | transferase activity, transferring phosphorus-containing groups (GO:0016772) | 0.44 | 2.84E-02 |
| La vs. Lc flower | none |  |  |
| La vs. Lc fruit | none |  |  |
| Lc flower vs. bud | pectinesterase activity (GO:0030599) | 10.69 | 1.48E-02 |
|  | RNA binding (GO:0003723) | 0.15 | 8.99E-03 |
| La flower vs. bud | sodium:proton antiporter activity (GO:0015385) | 14.17 | 2.35E-04 |
|  | cellulose synthase (UDP-forming) activity (GO:0016760) | 10.52 | 3.18E-02 |
|  | polygalacturonase activity (GO:0004650) | 8.13 | 8.50E-03 |
|  | iron ion binding (GO:0005506) | 3.12 | 3.48E-02 |
|  | oxidoreductase activity acting on paired donors with incorporation or reduction of molecular oxygen (GO:0016705) | 2.89 | 4.07E-02 |
|  | protein binding (GO:0005515) | 0.71 | 3.99E-02 |
|  | RNA binding (GO:0003723) | 0.17 | 1.30E-03 |
| Lc fruit vs. flower | heme binding (GO:0020037) | 5.11 | 9.54E-04 |
|  | hydrolase activity, hydrolyzing O-glycosyl compounds (GO:0004553) | 4.28 | 1.15E-03 |
| La fruit vs. flower | none |  |  |
| flower vs. bud | acid-amino acid ligase activity (GO:0016881) | 40.35 | 1.53E-02 |
| fruit vs. flower | none |  |  |

As we were interested in differences in the gene regulatory network involved in fruit dehiscence in the two species, known to be largely composed of transcription factors in *Arabidopsis thaliana* (reviewed in (Ballester and Ferrándiz 2017)), we analyzed genes annotated to have “DNA-binding transcription factor activity” (GO:0003700) in more detail. This set includes transcription factors and transcriptional regulators. For simplicity, we will refer to this dataset as genes encoding transcription factors (TFs).

When comparing flowers and buds, 21 and 28 TFs were DEGs in *L. campestre* and in *L. appelianum*, respectively. Among them, there are 13 TFs that were DEGs comparing these structures in both species, including four genes with known functions in flower development *AGAMOUS-LIKE 104* (*AGL104*) (Adamczyk and Fernandez 2009), *SPOROCYTELESS* (*SPL*, also termed *NOZZLE*) (Balasubramanian and Schneitz 2000), *ORESARA1* (*ORE1*, also termed *ANAC092, ATNAC2, ATNAC6*) (Gao et al. 2018) and *ZINC FINGER PROTEIN 2* (*ZFP2*) (Cai and Lashbrook 2008) (Table 4). Between fruits and flowers, there are 12 TFs in *L. campestre* and 23 TFs in *L. appelianum* that are DEGs. Five of these genes are DEGs in fruits versus flowers in both species (Table 4). TFs with differential expression between structures in both species are probably those TFs with common functions for flower and fruit development.

**Table 4:**
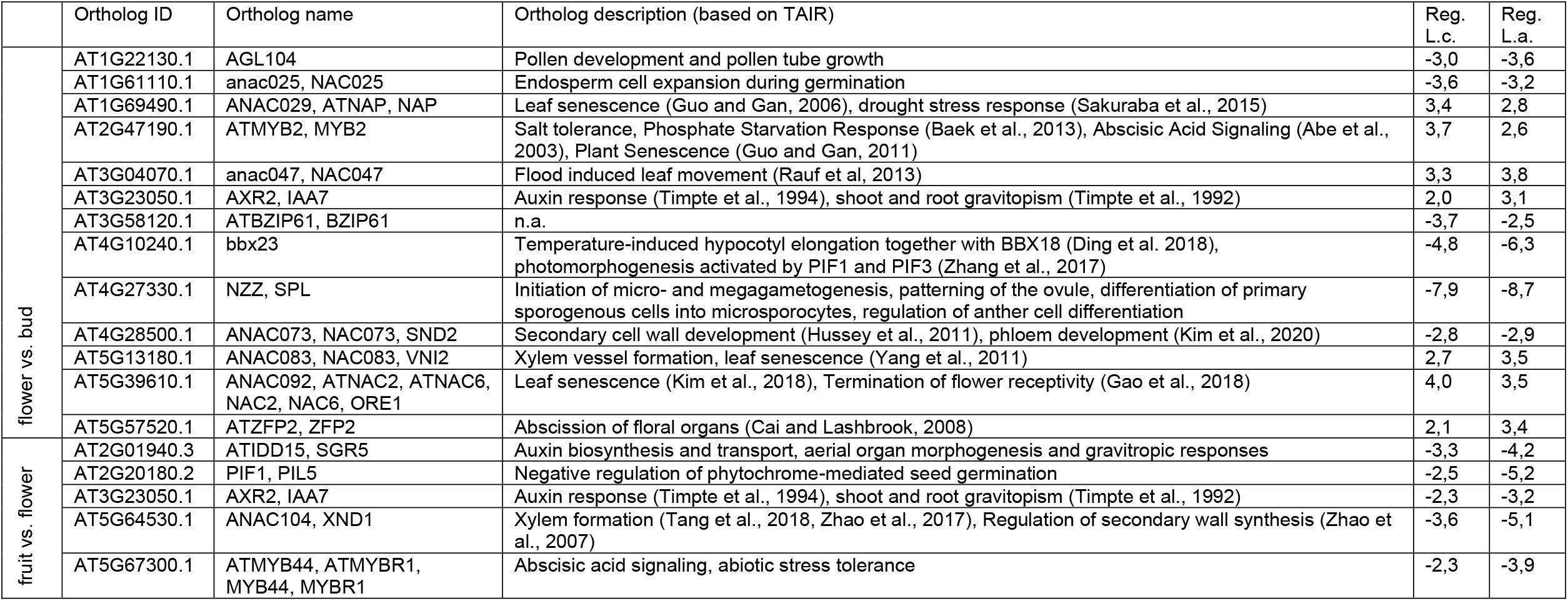
Differentially expressed genes in different structures annotated as “DNA-binding transcription factor activity” (GO:0003700).

When comparing the two species, 43 TFs were DEGs in buds, 68 in flowers and 49 in fruits. Among these TFs, 19 were DEGs in all structures (Table 5). Interestingly, four genes involved in flowering time determination, *SQUAMOSA PROMOTER BINDING PROTEIN-LIKE 4* (*SPL4*) (Jung et al. 2016), *NUCLEAR FACTOR Y-B2* (*NF-YB2*) (Cao et al. 2014), *NUCLEAR FACTOR Y-B10* (*NF-YB10*) (Wenkel et al. 2006) and *FLOWERING LOCUS C* (*FLC*) (Michaels and Amasino 1999), as well as the fruit development genes *FRUITFUL* (*FUL*) (Gu et al. 1998) and *YABBY3* (*YAB3*) (Dinneny et al. 2005; González-Reig et al. 2012) (Figure 5) were on this list.

**Table 5:** DEGs between the two *Lepidium* species annotated as “DNA-binding transcription factor activity”. (GO:0003700)

| Ortholog ID | Ortholog name | Ortholog description (based on TAIR) | Reg.<br>bud | Reg.<br>flower | Reg.<br>fruit |
| --- | --- | --- | --- | --- | --- |
| AT1G01060.1 | LHY | Involved in circadian rhythm | 2.5 | 2.4 | 2.8 |
| AT1G14687.1 | HB32, ZHD14 | n.a. | -4.0 | -3.1 | -2.2 |
| AT1G27370.1 | SPL10 | Development of lateral organs, lamina shape, lateral root growth (Yu et al., 2015) | -3.9 | -3.8 | -4.6 |
| AT1G46264.1 | HSFB4, SCZ | Asymmetry of stem cell divisions | -5.3 | -4.4 | -4.1 |
| AT1G53160.2 | FTM6, SPL4 | Regulation of flowering and vegetative phase change | -5.2 | -5.3 | -4.5 |
| AT1G79840.2 | GL2 | Regulation of epidermal cell identity, regulation of seed oil content | 3.5 | 3.6 | 2.7 |
| AT3G09370.2 | MYB3R-3 | DNA damage response | -2.4 | -2.2 | -2.6 |
| AT3G11280.1 | n.a. | n.a. | -2.6 | -2.9 | -2.9 |
| AT3G14020.1 | NF-YA6 | Involved in male gametogenesis, embryogenesis, and seed development (Mu et al, 2012) | -2.6 | -2.2 | -2.4 |
| AT3G53340.1 | NF-YB10 | Flowering time determination (Tao et al., 2017) | 3.3 | 3.4 | 4.9 |
| AT4G00180.1 | YAB3 | Specification of abaxial cell fate, involved in fruit patterning along with FIL | -2.6 | -2.5 | -2.5 |
| AT4G01280.2 | RVE5 | Clock regulation, growth regulation (Gray et al., 2017) | 2.6 | 2.2 | 2.2 |
| AT4G31060.1 | n.a. | n.a. | -2.9 | -3.5 | -3.4 |
| AT5G04340.1 | C2H2, CZF2, ZAT6 | Phosphate homeostasis (Devaiah et al., 2007), Cd accumulation and tolerance (Chen et al. 2016) | 4.0 | 4.6 | 3.4 |
| AT5G10140.1 | AGL25, FLC, FLF, RSB6 | Flowering time determination | 3.7 | 4.1 | 5.6 |
| AT5G39760.1 | HB23, ZHD10 | Light-induced development (Perrella et al., 2018) | 7.2 | 6.3 | 4.9 |
| AT5G41920.1 | SCL23 | Endodermis development (Yoon et al., 2016; Cui et al., 2014) | -5.0 | -5.3 | -3.9 |
| AT5G47640.1 | NF-YB2 | Flowering time determination (Hou et al., 2014) | 2.2 | 2.6 | 2.4 |
| AT5G60910.1 | AGL8, FUL | Fruit development (Gu et al, 1998), apical hook development (Führer et al., 2020) | -2.9 | -2.4 | -2.7 |

**Figure 5:**
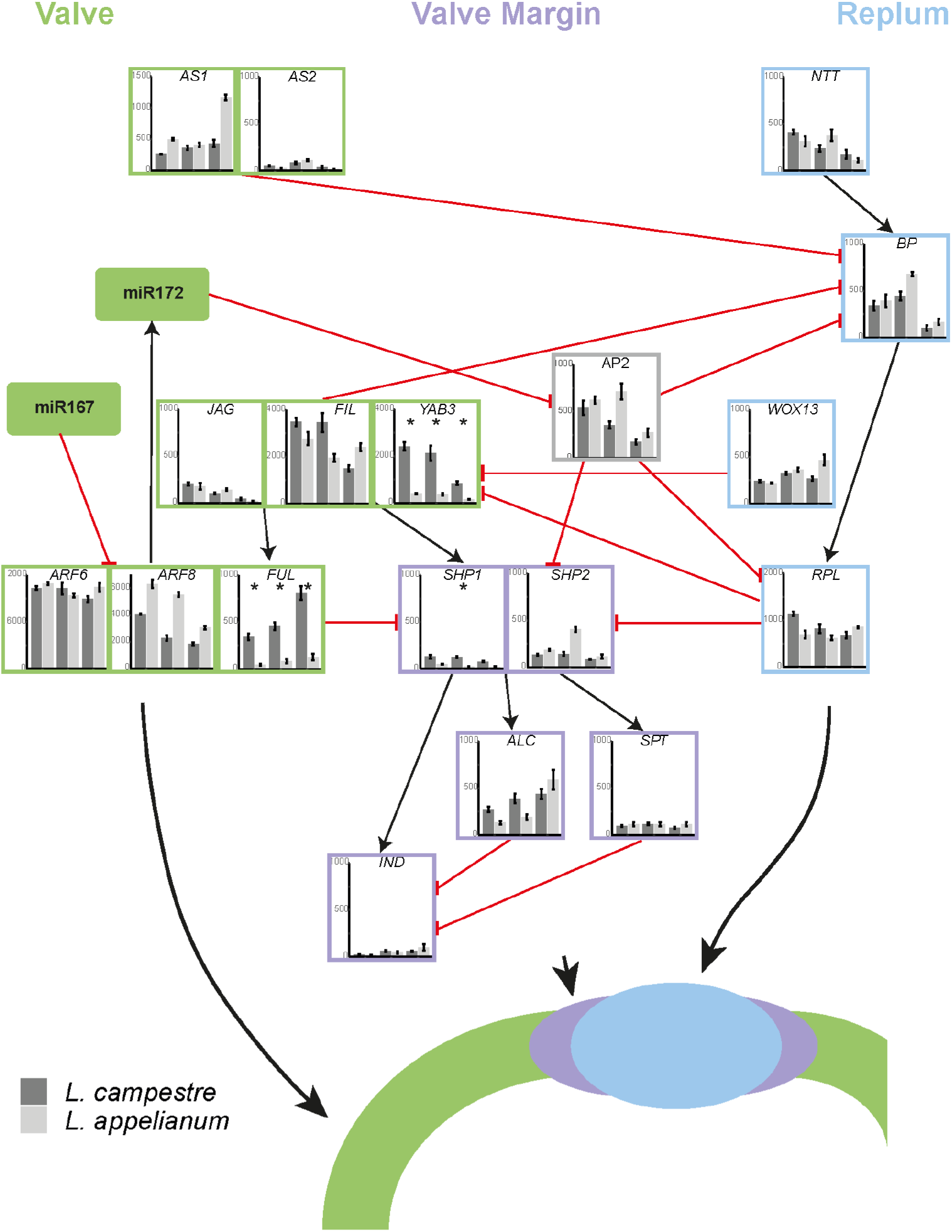
Gene regulatory network for the development of valve, valve margin and replum of a fruit. The network is based on what has been determined in *A. thaliana* and is modified after Chavez-Montez *et al*., 2015. Relative expression levels of genes in *L. campestre* and *L. appelianum* as determined in this study by transcriptome analysis are shown. Significant differences between *L. appelianum* and *L. campestre* are indicated by asterisks (P ≤0.05).

Two TFs were found to be DDEGs when comparing flowers and buds in the two species (Table 6), among them *MASSUGU 2* (*MSG2*, also known as *INDOLE-3-ACETIC ACID INDUCIBLE 19*) (Tatematsu et al. 2004) which has been shown to be involved in stamen filaments development (Tashiro et al. 2009). Comparing fruits and flowers, seven TFs, *PHY-INTERACTING FACTOR 1* (*PIF1*, also known as *PHYTOCHROME INTERACTING FACTOR 3-LIKE 5*) (Huq et al. 2004), *MYB DOMAIN PROTEIN 57* (*MYB57*) (Bender et al. 2013), *TCP FAMILY TRANSCRIPTION FACTOR 4* (*TCP4*, also known as *MATERNAL EFFECT EMBRYO ARREST 35*) (Nag et al. 2009), *BRANCHED 1* (*BRC1*, also known as *TCP FAMILY TRANSCRIPTION FACTOR 18*) (Aguilar-Martínez et al. 2007), *REVEILLE 6* (*RVE6*) (Hsu et al. 2013), *TRIPTYCHON* (*TRY*) (Schnittger et al. 1999) and *OBF BINDING PROTEIN 4* (*OBP4*, also termed *DOF5*.*4*) (Xu et al. 2016) are DDEGs in *L. appelianum* as compared to *L. campestre*.

**Table 6:** DDEGs in different structures annotated as “DNA-binding transcription factor activity” (GO:0003700)

|  | Ortholog ID | Ortholog name | Ortholog description (based on TAIR) | Reg. |
| --- | --- | --- | --- | --- |
| flower<br>vs. bud | AT3G15540.1 | IAA19, MSG2 | Stamen filaments development | 4,8 |
|  | AT5G47230.1 | AtMACD1, ERF102, ERF5 | Stress response, leaf growth | 5,3 |
| fruit vs. flower | AT2G20180.2 | PIF1, PIL5 | Phytochrome-mediated seed germination | 2,7 |
|  | AT3G01530.1 | ATMYB57, MYB57 | Stamen and nectary development (Bender et al., 2013) | -2,9 |
|  | AT3G15030.1 | MEE35, TCP4 | Cotyledon, leaf and petal development, seed oil accumulation<br>Overexpression of TCP4 causes increased deposition of lignin and cellulose (Sun et al., 2017), TCP4 interacts with AS2 to regulate BP? (Li et al., 2012) | -4,3 |
|  | AT3G18550.1 | BRC1, TCP18 | Arrests axillary bud development and prevents axillary bud outgrowth. Role in flowering control. | -4,3 |
|  | AT5G52660.2 | RVE6 | Involved in circadian rhythm | -2,6 |
|  | AT5G53200.1 | TRY | Trichome and root hair patterning, phosphate starvation response | -6,0 |
|  | AT5G60850.1 | DOF5.4, OBP4 | Cell Cycle Progression and Cell Expansion | -2,3 |

### Extension of the gene regulatory network for fruit development

We next investigated how the TFs shown to be differentially regulated between fruits and flowers may be involved in the gene regulatory network controlling fruit development (Figure 5). Therefore, we searched for binding sites of the seven TFs identified by chromatin immunoprecipitation followed by sequencing (ChIP-seq) experiments in the promotors of the genes known to be involved in fruit development. On ChIP-Hub (Chen et al. 2019), no ChIP-seq data is available for BRC1 and for TRY.

Binding of OBP4 was found in the promotor of all but one of the 18 fruit development genes (Table 7). Binding of RVE6, MYB57, PIF1 and TCP4 was detected in the promotors of 11, 7, 5 and 2 fruit development genes, respectively. PIF1 predominantly binds to the promotors of valve identity genes, with binding to four out of eight valve identity gene promotors and apart from that only binding to one of five valve margin genes. *ARF8* is the only fruit development gene for which none of the differentially regulated genes was found to bind to its promotor. To the promotors of *YAB3* and *FUL*, which were found to be differentially expressed in all structures between *L. campestre* and *L. appelianum*, binding of TCP4, RVE6 and OBP4 and of MYB57, RVE6 and OBP4, respectively, was found.

**Table 7:**
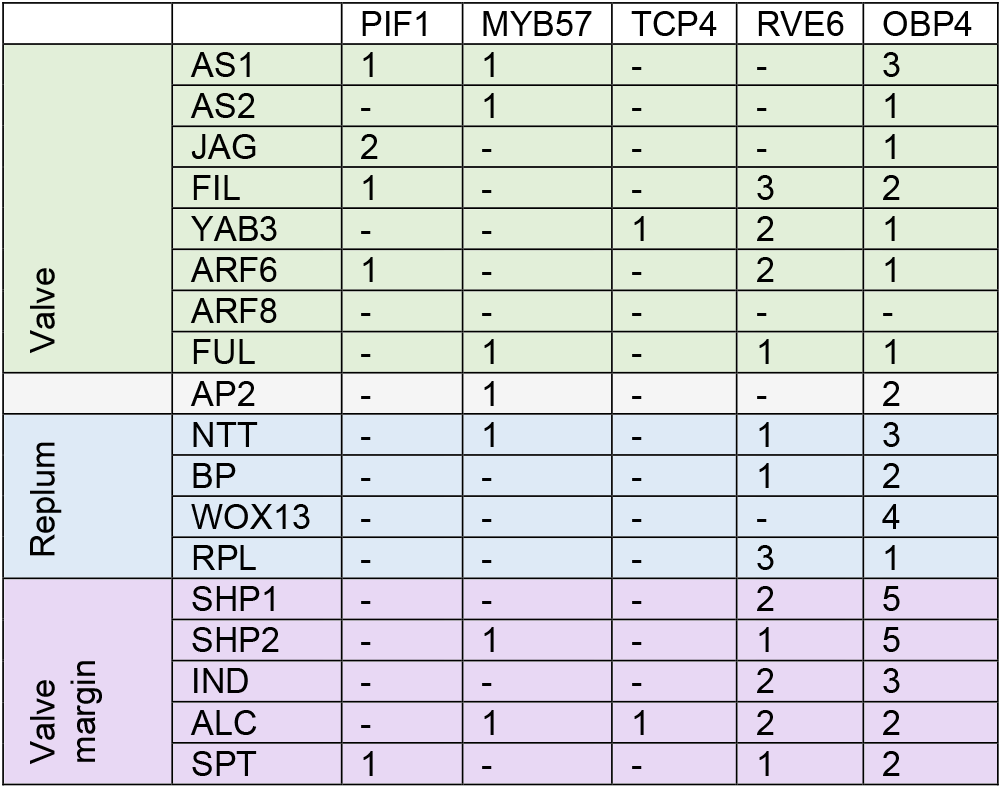
Binding of TFs found to be DDEGs to the promotors of known fruit development genes.

## Discussion

### *Transcriptomes and small RNA datasets of L. campestre and L. appelianum* are nearly complete

We have sequenced the transcriptomes of floral buds, flowers and fruits of *L. campestre* and *L. appelianum*. Benchmarking of Universal Single-Copy Orthologs (BUSCO) analysis revealed that the transcriptome assemblies of the two species contain more than 94% of the eudicotyledonous “near-universal single-copy orthologs”. This number is similar to or more than that for transcriptome assemblies of other Brassicaceae (Chandler et al. 2020; Yao et al. 2020; Fernandez-Pozo et al. 2021). Furthermore, we found members of all 16 miRNA families that were found to have originated before the emergence of eudicots and conserved in eudicots (Montes et al. 2014). These findings reveal that our transcriptome and small RNA data includes most of the expected transcripts and miRNAs.

### Differences in gene expression mainly between floral structures

We identified more DEGs when comparing the same structure between the two species than comparing different structures in the same species (Figure 4). This indicates that gene regulation has diverged between the two species. This is different to what has been observed other flowering plant species, where the correlation of gene expression is higher in the same structure of different species than in different structures of the same species (Chanderbali et al. 2010). However, in this case microarray expression data was analyzed which may select for conserved genes and structures at similar developmental time points were compared.

The highest number of DEGs was observed in flowers, followed by fruits and buds, and the highest number of DDEGs was found in fruits versus flowers as compared to flowers versus floral buds. This indicates that the differences in gene expression between *L. campestre* and *L. appelianum* are most pronounced between flowers and in the transition from flowers to fruits. This is expected as the developmental program leading to fruit dehiscence or indehiscence needs to be initiated before the fruits are formed. Supportingly, in *Aethionema arabicum*, a plant that develops dehiscent and indehiscent fruits on the very same individual, differences between the fruit types start to occur in flowers after anthesis (Lenser et al. 2018).

A number of GO terms related to cell wall synthesis and degradation, e.g. pectinesterase activity, cellulose synthase activity and polygalacturonase activity were overrepresented in different sets of DEGs. It has been recognized that secondary cell wall formation at the valve margins (Mitsuda and Ohme-Takagi 2008) and degeneration of cell walls in the separation layer are essential processes for fruit dehiscence after the DZ is correctly specified (Ogawa et al. 2009). Hence, the overrepresentation of GO terms related to cell wall synthesis and degradation is not surprising.

### Confirmation of previous expression study

In a previous study, we have compared expression of the valve margin genes as well as the valve gene *FUL* and the replum gene *RPL* between *L. campestre* and *L. appelianum* by *in situ* hybridization (Mühlhausen et al. 2013). We showed that their orthologues from *L. campestre* (dehiscent fruits) are similarly expressed as in *A. thaliana* while expression of the respective orthologues is abolished in valve margins of indehiscent *L. appelianum* fruits. Analysis using qRT-PCR revealed that the valve margin genes *IND* and *SHP1* are expressed at a significantly higher level in flowers and early fruits (the fruit stage for which the transcriptome was sequenced here) of *L. campestre* than in *L. appelianum*. Significantly higher expression was confirmed in the present study for *SHP1* in flowers (Figure 5). qRT-PCR analysis revealed significantly higher expression of *SHP2* in flowers and of *ALC* in early fruits in *L. appelianum*. Expression was not significantly different in the present transcriptome analysis, but expression was also found to be higher for *SHP2* in flowers and for *ALC* in fruits. Like qRT-PCR analysis, our transcriptome analysis also found significantly higher expression of *FUL* in fruits of *L. campestre*. Similarly, *RPL* was found to be expressed at a higher level in the flowers of *L. campestre* though the difference was only found to be significant in qRT-PCR but not in transcriptome analysis. *AP2* was found to be expressed at a lower level in flowers and fruits of *L. campestre* by both analyses. Again, the difference was significant in qRT-PCR analysis but not in transcriptome analysis. Hence, our transcriptome analysis is in good agreement with the previous qRT-PCR analyses, but differences were less often significant.

### Known flower and fruit development genes are differentially expressed

To identify differences in the regulation of flower and fruit development between *L. campestre* and *L. appelianum*, we focused on differentially expressed or differentially differentially expressed transcription factors (TFs). Among 19 TFs which were found to be differentially expressed in all three examined structures, four TFs are involved in flowering time determination. *L. appelianum* and *L. campestre* have different flowering periods according to the Jepson Herbarium (Jepson Flora Project (eds.) 2021, Jepson eFlora, https://ucjeps.berkeley.edu/eflora/, accessed on May 25, 2021), which may be caused by differences in the expression of the identified flowering time genes.

As mentioned above, the fruit development genes *SHP1* and *FUL* were found to be differentially expressed. *FUL* expressed at a significantly lower level in *L. appelianum* in all three structures. In *A. thaliana*, the *ful* knockout mutation causes indehiscence in (Gu et al. 1998; Ferrandiz et al. 2000). Furthermore, *YAB3* (Dinneny et al. 2005; González-Reig et al. 2012) (Figure 5) was also differentially expressed in floral buds, flowers and fruits. FUL is activated by *JAG, FIL* and *YAB3* (Dinneny et al. 2005). Concordantly, we found that *YAB3* has a significant higher expression level in *L. campestre* than in *L. appelianum*. Expression of *JAG* and *FIL* is not significantly different between both species. *yab3* single mutants do not have any major defects in dehiscence but *fil yab3* double mutants are largely indehiscent (Dinneny et al. 2005). Hence, decreased expression of *YAB3* in *L. appelianum* as compared to *L. campestre* may have been an important factor for the evolutionary shift from dehiscent to indehiscent fruits in *L. appelianum*. This also shows that there was not only a change in the control of valve margin identity genes but also of the valve identity genes and shifts the causative mutation further upstream in the gene regulatory network of fruit development.

### MiR166 is differentially regulated in fruits versus flowers

Our smallRNA sequencing revealed that the miRNA homologous to miR165a-3p, miR165b, miR166a-3p, miR166b-3p, miR166c, miR166d, miR166e-3p, miR166f and miR166g (Reinhart et al. 2002) is encoded by a DDEG when comparing fruits and flowers. Targets of miR165 and miR166 are the mRNAs of HD-Zip transcription factors like PHABULOSA (PHB), REVOLUTA and PHAVOLUTA (Rhoades et al. 2002). Recently, a function of the miR166-PHB module in anther dehiscence has been elucidated (Li et al. 2019). Upregulation of miR166 in the *jba-1D* mutant leads to downregulation of its target gene PHB which results in increased expression of *SPOROCYTELESS/NOZZLE* (*SPL/NZZ*). *jba-1D* mutants do not develop a dehiscence zone in anthers, i.e. overexpression of miR166 leads to indehiscence of anthers. Expression of miR166 in fruits is much higher in *L. appelianum* (indehiscent fruits) than in *L. campestre* (dehiscent fruits), while the opposite is the case in flowers (Figure 6). Hence, miR166 may have a role in the development of indehiscent fruits in *L. appelianum* though the details of the regulation remain to be elucidated.

**Figure 6:**
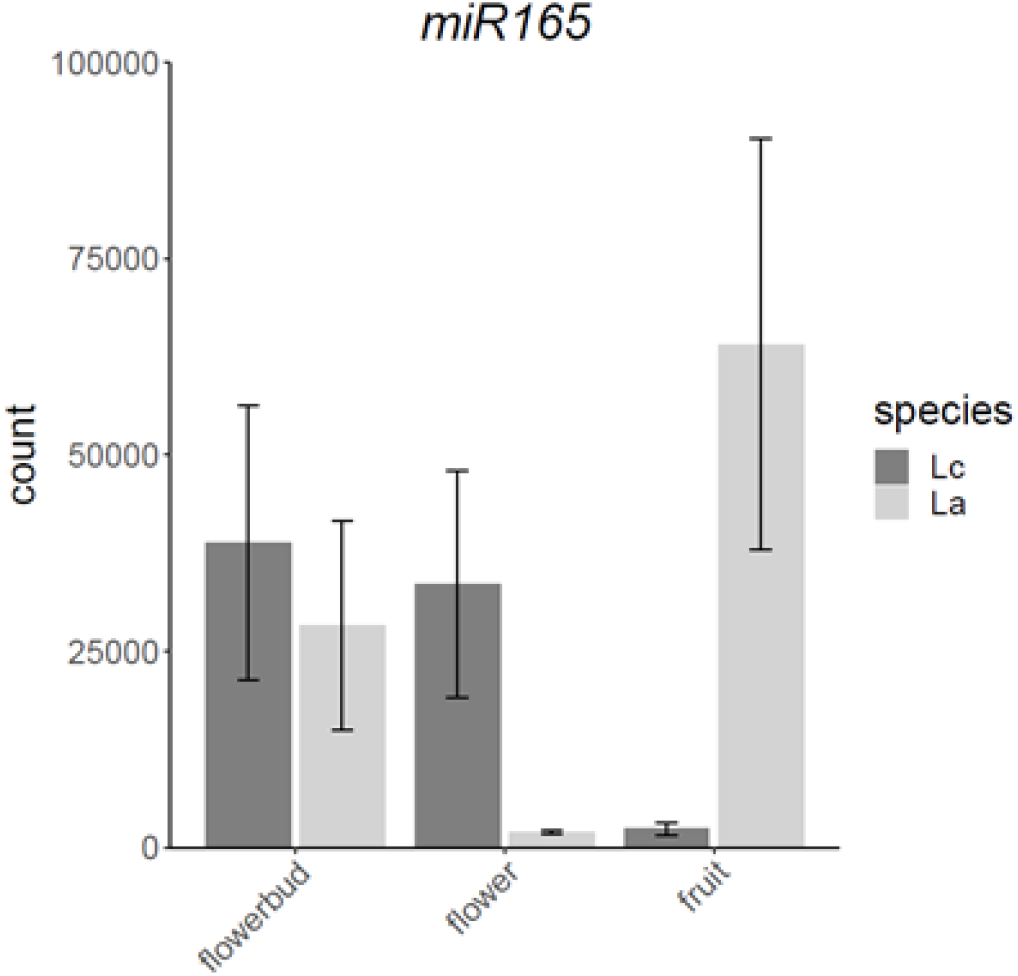
Expression data plot of the miRNA homologous to miR165a-3p of *A. thaliana*. miR165a-3p is identical to miR165b, miR166a-3p, miR166b-3p, miR166c, miR166d, miR166e-3p, miR166f and miR166g such that they cannot be distinguished and hence are summarized here as miR165. Bars indicate mean normalized count values of reads mapping to miR165 in the corresponding structure and species. Dark and light grey bars represent the mean values for *L. campestre* (Lc) and for *L. appelianum* (La), respectively. The error bars indicate the standard deviation.

### BRC1 *and* TCP4 *as candidate genes for the evolutionary shift from dehiscent to indehiscent fruits*

Our transcriptome analysis also identified seven TFs belonging to DDEGs when comparing flowers and fruits (Table 6). PIF1 is a basic helix-loop-helix (bHLH) transcription factor that negatively regulates chlorophyll biosynthesis (Huq et al. 2004); it is involved in a variety of biological processes such as the repression of light-induced seed germination and chlorophyll accumulation in light (Castillon et al. 2007). RVE6 is a MYB gene that controls the pace of the circadian clock together with its close homologs RVE4 and RVE8 (Hsu et al. 2013). The zinc finger gene OBP4 functions in cell cycle progression and cell expansion (Xu et al. 2016) and is involved in root development (Rymen et al. 2017; Ramirez-Parra et al. 2017). So far, involvement of these three factors in flower and fruit development has, to the best of our knowledge, not been reported.

Two other MYB genes, MYB57 and TRY have also been found to be differentially regulated (Table 6). MYB57 functions redundantly with MYB21 and MYB24 to regulate stamen development (Cheng et al. 2009). TRY controls the spacing pattern of trichomes, which are single-celled hairs (Schnittger et al. 1999). Recently it has been found that TRY and other MYB genes of the regulatory network for trichome patterning have been modulated to trigger trichome development in fruits (Arteaga et al. 2021). Hence, these two gene are known to function during flower and fruit development but association with fruit dehiscence is not known so far.

More interestingly, the genes encoding for the two TCP transcription factors BRC1 and TCP4 are DDEGs between fruits and flowers when comparing *L. campestre* and *L. appelianum*. Expression of *BRC1* correlates with bud inhibition (Aguilar-Martínez et al. 2007; Braun et al. 2012) but recently, it has been shown that *BRC1* is neither necessary nor sufficient for bud inhibition (Seale et al. 2017). Noticeably, it has been hypothesized that *BRC1* may guide fruit morph determination in the dimorphic Brassicaceae plant *Aethionema arabicum* (Lenser et al. 2018). *Ae. arabicum* produces two fruit morphs on the same plant, one of which is dehiscent and the other one is indehiscent. qRT-PCR analyses showed that the expression of *BRC1* in *Ae. arabicum* is high in flowers and decreases strongly in the indehiscent morph but remains at a low level in flowers and fruits of the dehiscent morph. We observe a very similar pattern in our transcriptome analysis for the indehiscent morph *in L. appelianum* and the dehiscent morph in *L. campestre* (Figure 7). It is not known, however, as to how *BRC1* interacts with the genes of the fruit development network (Figure 5).

**Figure 7:**
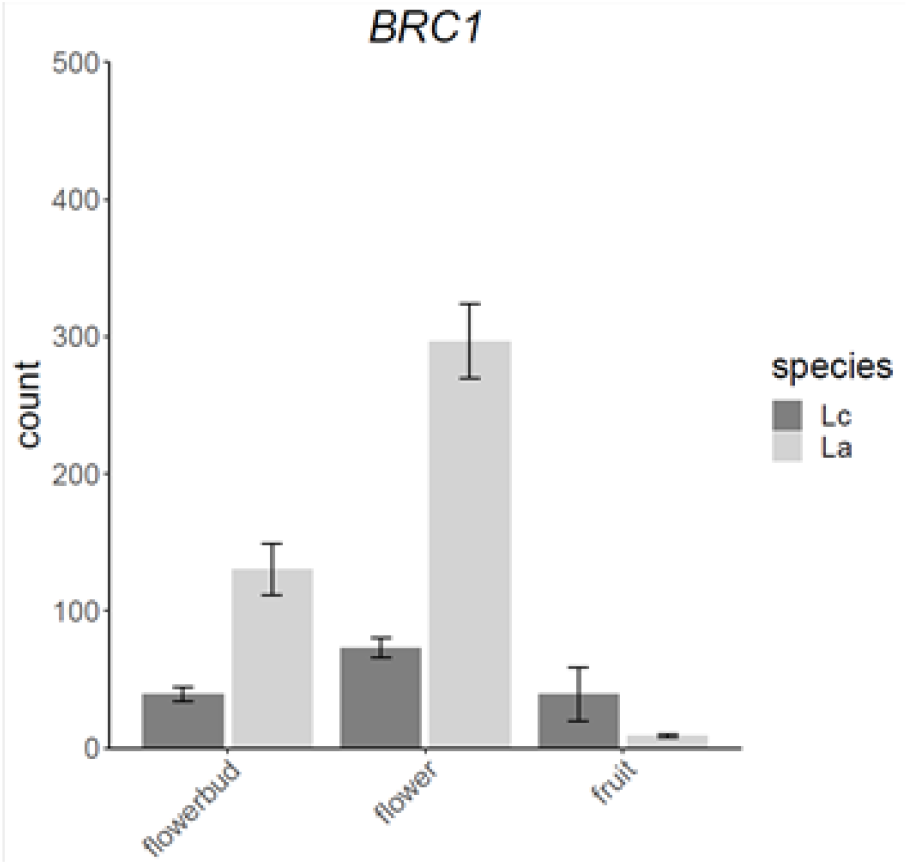
Expression data plot of *BRC1* in *L. campestre* (Lc) and *L. appelianum* (La). Bars indicate mean normalized count values of reads mapping to *BRC1* in the corresponding structure and species. Dark and light grey bars represent the mean values for *L. campestre* (Lc) and for *L. appelianum* (La), respectively. The error bars indicate the standard deviation.

TCP4 has been found to be involved in leaf and flower development as well as in seed oil biosynthesis in *A. thaliana* (Kong et al. 2020; Nag et al. 2009; Palatnik et al. 2003). Furthermore, TCP4 directly activates the expression of *miR167* which targets the TFs ARF6 and ARF8 (Rubio-Somoza and Weigel 2013). This regulation has been hypothesized to be important for flower maturation, but may also be involved in fruit dehiscence as ARF6 and ARF8 are part of the gene regulatory network of fruit development (Zheng et al. 2019) (Figure 5). Another study found physical interaction of TCP4 and AS2 in yeast-two-hybrid experiments (Li et al. 2012). AS2 has also previously been found to be involved in fruit patterning (Alonso-Cantabrana et al. 2007) (Figure 5). Our analysis of ChIP-seq data on ChIP-Hub (Chen et al. 2019) additionally revealed that TCP4 binds to the promotor of *YAB3* (Table 7), which has been found to be differentially expressed between *L. campestre* and *L. appelianum* in all structures examined. In flowers, TCP4 is expressed at a higher level in *L. appelianum* than in *L. campestre* while the expression pattern is the other way round for *YAB3*. Hence, it is conceivable that TCP4 represses *YAB3* in flowers.

## Conclusions

Taken together, our study provides insights into the gene regulatory differences in fruit development between *L. campestre* producing dehiscent fruits and *L. appelianum* forming indehiscent fruits. We confirm differences in the expression of the fruit development genes *SHP2* and *FUL* between the two species and reveal the importance of the valve identity gene *YAB3* for fruit indehiscence in *L. appelianum*. We uncover the microRNA miR166 and the TCP transcription factors BRC1 and TCP4 as new candidates for causing the evolutionary transition from dehiscent to indehiscent fruits in *L. appelianum*.

## Methods

### Plant material, RNA extraction and sequencing

Seeds were sawn on a mixture of seedling substrate (Kammlott, Kammlott GmbH, Erfurt, Germany)/sand/vermiculite (1–3 mm) (8:1:1) which was supplemented with g ™ each of Osmocote mini (www.scotts.com,The Scotts Miracle-Gro Company, Marysville, OH, USA) and Triabon (http://www.compo-expert.com, COMPO Expert GmbH, Münster, Germany). The seeds were placed for 4 days at 4°C for stratification and then put in the greenhouse under a light-dark cycle of 16 hours light, 8 hours dark of artificial light, plus daylight. After 5 weeks in the greenhouse, the plants were vernalized for at least 13 weeks at 4°C with a light-dark cycle of 8 hours light, 16 hours dark. After vernalization, the plants were put back in the greenhouse. Plant material was harvested from two batches of independently grown plants 3 to 5 weeks after the end of vernalization. Plant material was harvested between 12pm and 4pm to minimize the effect of circadian rhythm. Late flower buds, flowers and early fruits were harvested and immediately frozen in liquid nitrogen. Three samples were taken for each species and each structure resulting in 18 samples in total. For each sample, about 100mg plant material was pooled from four individual plants. The material was pulverized in liquid nitrogen using a mortar and pestle. RNA was extracted using Qiazol (Qiagen) according to the manufacturer’s instructions. RNA quantity and quality was checked by gel electrophoresis. The samples were sent to the Vienna BioCenter Facility for Next Generation Sequencing where they were quality checked and sequenced on a HiSeqV4. For mRNA sequencing, 125bp paired-end reads were produced and 50bp single-end reads were generated for small RNA sequencing.

### Preprocessing of RNA-seq data

Raw reads were corrected using Rcorrector (Song and Florea 2015) with default settings. Uncorrectable reads were excluded using a python script obtained from https://informatics.fas.harvard.edu/best-practices-for-de-novo-transcriptome-assembly-with-trinity.html which was slightly modified for excluding uncorrectable reads from smallRNA libraries. The remaining reads were trimmed with Trim Galore! (https://www.bioinformatics.babraham.ac.uk/projects/trim_galore/) using the following settings: `--clip_R1 12, --clip R2 12, --paired, --retain_unpaired, --phred33, --length 36, -q 5, --stringency 5, –e 0.1` for transcriptome reads and the following settings --phred33, --length 18, --max_length 26, -q 5, --stringency 5, –e 0.1, -a *adapter* for small RNA reads where `adapter` was replaced by the corresponding adapter sequence identified using FastQC (https://www.bioinformatics.babraham.ac.uk/projects/fastqc/). Thereafter, Poly-A and Poly-T tails were removed from transcriptome reads with PrinSeq (Schmieder and Edwards 2011) using the settings -`trim_tail_left 5` and `-trim_tail_right 5`. Reads that mapped to the genome of *Frankliniella occidentalis* (GenBank: GCF_000697945) or to rRNAs (GenBank: X52320.1), mitochondrial (GenBank: Y08501.2) or chloroplast (GenBank: AP000423.1) sequences from *A. thaliana* as determined using bowtie2 (settings: `--very-sensitive-local, --phred33`) (Langmead and Salzberg 2012) were excluded from further analyses from both, the transcriptome and the smallRNA libraries.

### De novo assembly

To simplify *de novo* assembly, also duplicate reads, i.e. reads with the exact same length and sequence, were removed. The remaining reads were assembled using Trinity (Grabherr et al. 2011b) with default settings for the two species *L. campestre* and *L. appelianum* separately. To identify remaining contamination in the transcriptome, a BLASTn search was conducted against the nucleotide sequence database of NCBI (nt) using the transcripts in the assembly as query with the option `-max_target_seqs 5`. Transcripts for which the best BLASTn result came from a non-plant species and had an eValue of E<10^−10^ were removed from the transcriptomes. The completeness of the assembled transcriptomes was evaluated using the Benchmarking Universal Single-Copy Orthologs tool BUSCO (Simao et al. 2015) and their accompanying dataset for eudicotyledons with 2121 groups (odb10).

### Separation of chimeras in the assemblies

The initial assemblies contained chimeras composed from two different transcripts. As these chimeras often result from misassembly (Yang and Smith 2013a), we sought to separate chimeras into their separate transcripts. To recognize chimeras, we first conducted a BLASTn search (Altschul et al. 1990) using the transcripts from the *Lepidium* transcriptomes as query and the cDNA sequences of the representative *A. thaliana* gene model as provided by TAIR10 (TAIR10_cdna_20110103_representative_gene_model_updated.fasta) as database saving the best two subjects (i.e. *A. thaliana* cDNAs) for each query (i.e. each transcript from the *Lepidium* transcriptomes) (Figure 8). Using a customized perl-script (Supplementary Data S1) we searched for instances where the two subjects fitted to different regions of the query (i.e. one part of the *Lepidium* transcript has a BLAST hit corresponding to one *A. thaliana* cDNA while another part of the same transcript has a BLAST hit corresponding to another *A. thaliana* cDNA). These instances likely indicate chimeric *Lepidium* transcripts. To identify the best position to split the chimeras, we considered at which position and to what extend the *A. thaliana* cDNAs matched to the *Lepidium* transcripts as shown in Figure 8. Chimeras were split if the overlap was less than 150 nucleotides either in the middle of the overlap or at the positions corresponding to the corrected end and beginning of the involved transcripts (Figure 8).

**Figure 8:**
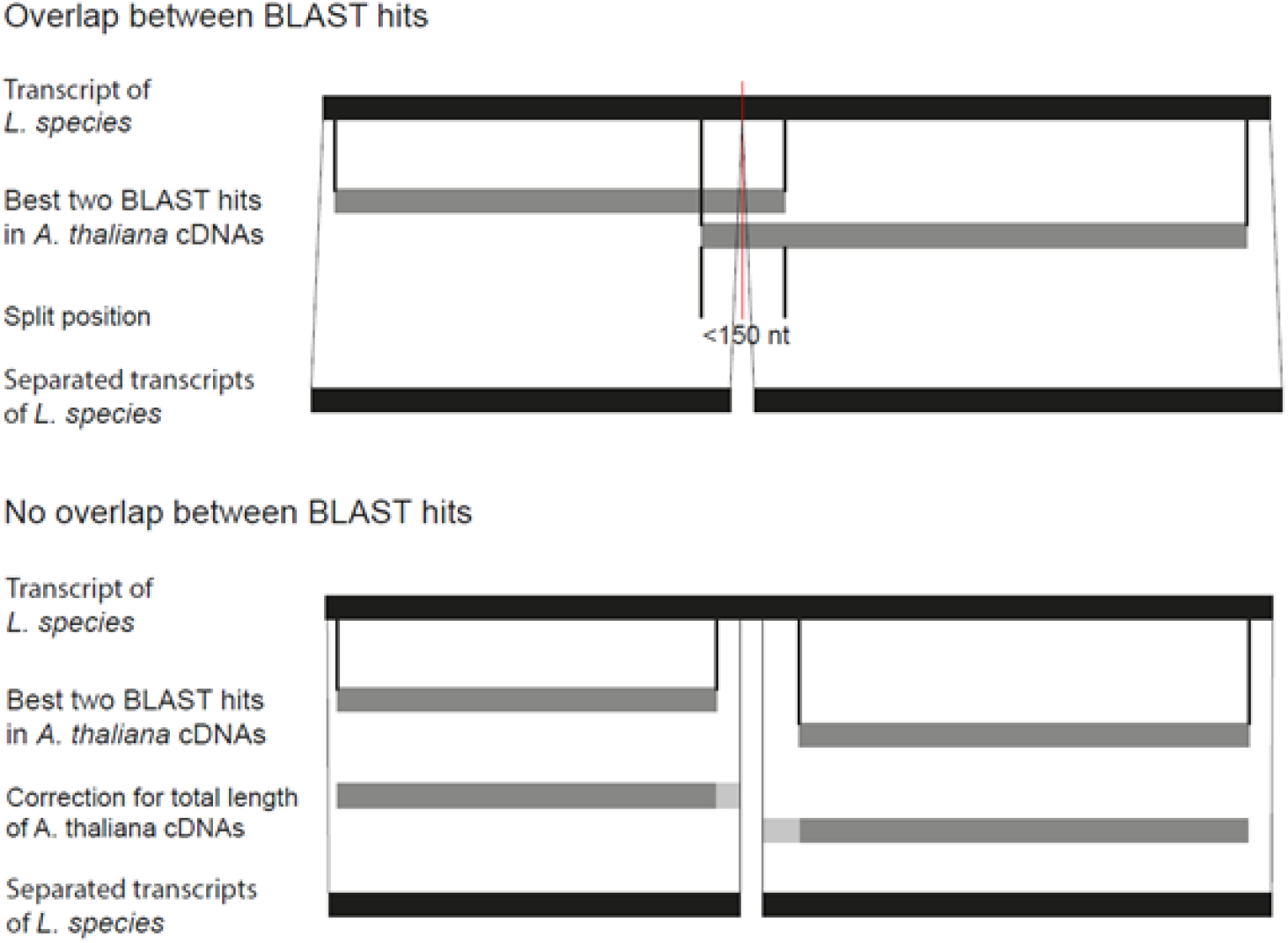
Schematic presentation of the detection and separation of chimeric transcripts in the *Lepidium* transcriptomes. The procedure is slightly different depending on whether the positions of the best two BLAST hits in *A. thaliana* cDNAs overlap on the *Lepidium* transcript or not. If the positions of the best two BLAST hits overlap by less than 150 nucleotides, the *Lepidium* transcript is split in the middle of the overlap. Otherwise, the beginning and end of the involved transcripts was determined based on the total length of the fitting *A. thaliana* cDNAs.

### Identification of miRNAs in smallRNA libraries

SmallRNA reads were mapped to mature miRNAs from *Arabidopsis thaliana* as downloaded from miRBase (Kozomara et al. 2019b) employing bowtie2 with the settings `-N 1, -L 18`. If a read mapped to a specific miRNA from *A. thaliana* this miRNA was considered to be present in the corresponding *Lepidium* species. Mature miRNAs only differing by one nucleotide were combined to avoid multiple mapping during read-counting. To identify novel miRNAs in *Lepidium*, we used ShortStack (Johnson et al. 2016) with the parameters `–foldsize 500,` `--dicermin 18` and the trinity transcriptome assembly of the corresponding *Lepidium* species as reference “genome”. For *L. campestre*, ShortStack was run a second time, this time using the genome sequence of *L. campestre* as available at the National Centre for Biotechnology Information (Sayers et al. 2020) as reference. The stem-loop sequences classified as “N” or “Y” by ShortStack were used as uery se uences for B AST searches of pre-miRNAs of *A. thaliana* as downloaded from miRBase (Kozomara et al. 2019b) to distinguish known from novel miRNAs. The stem-loop sequences classified as “Y” by ShortStack without similarity to pre-miRNAs of *A. thaliana* were classified as novel *Lepidium* miRNAs.

### Determination of orthologs

For transcriptome data, putative ortholog pairs were determined using the reciprocal best hit approach as follows. BLASTn searches were conducted using the transcriptome assembly with chimeras separated of *L. campestre* as query and the transcriptome assembly with chimeras separated of *L. appelianum* as subject and *vice versa*. For each transcript of *L. campestre* the best BLAST hits (having the same eValue, score and alignment length) in *L. appelianum* were recorded and *vice versa*. If a transcript T_c_ from *L. campestre* had the transcript T_a_ from *L. appelianum* among its best BLAST hits and transcript T_a_ from *L. appelianum* had transcript T_c_ from *L. campestre* in its list of best BLAST hits, these were considered as best reciprocal BLAST hit. Best reciprocal BLAST hits with an alignment length of more than 250 nucleotides and where the length of the shorter sequence was at least 50% of that of the longer sequence were considered as putative ortholog pairs. Additionally, another BLASTn search was conducted using the transcripts in the transcriptomes as query and the *Arabidopsis thaliana* TAIR10 cDNA dataset as database. The set of putative orthologous transcript pairs was pruned such that only one transcript isoform was kept for each species unless different isoforms fitted to different *A. thaliana* genes. The isoform with the longest alignment length between the two species was chosen to be kept. This way, for each transcript in the one transcriptome exactly one transcript in the other transcriptome was kept. We refer to this dataset as the ortholog-transcriptome. The transcripts in the ortholog-transcriptome dataset were named using the TAIR10 identifier of the best BLAST result or numbered if no BLAST result was obtained this way.

For the miRNA data, orthologous miRNAs of the two *Lepidium* species were defined as those miRNAs fitting to the same miRNA from *A. thaliana*. Comparison of the novel *Lepidium* miRNAs revealed that none of these was found in both species.

### Read mapping and feature counting

Preprocessed transcriptome and small RNA reads were mapped against ortholog-transcriptome and mature miRNAs, respectively, using bowtie2 (settings: `--very-sensitive-local, --phred33` and settings: `--phred33, -N 1, -L 18`, respectively) (Langmead and Salzberg 2012). A custom GFF was generated with one feature for each transcript and miRNA. Mapped reads per feature were then counted using HTSeq-count (Anders et al. 2015a) with the settings `–s no –t transcript –m union`.

### Differential gene expression analysis pipeline

Differentially expressed genes were identified using R (https://www.r-project.org/) and the Bioconductor packages edgeR (Robinson et al. 2010b) and DESeq2 (Love et al. 2014b). Transcript counts were normalized with respect to transcript length. Lowly expressed transcripts with normalized counts and lowly expressed miRNAs with raw counts of less than 19 were discarded. Considering the two species *L. campestre* and *L appelianum* and the structures bud, flower and fruit, the following multi-factor design was used: `species + structure + species:structure`. A Likelihood Ratio Test (LRT) and a quasi-likelihood F-test were conducted in DESeq2 (command: `DESeq(object, test=“LRT”, reduced=∼species + structure)`) and EdgeR (command: `glmQLFit(object,` `design)`), respectively to identify differentially expressed and differently differentially expressed genes. Only transcripts and miRNAs having a log-fold change to the base of 2 of more than 1 were considered. For DESeq2 the false discovery rate threshold α was set to .001. For principal component analysis, count data was normalized using regularized logarithm with the option `blind=FALSE` in DESeq2 and the principal components were plotted using the plotPCA function in R.

### GO enrichment analysis

Gene Ontology (GO) enrichment analysis was conducted on the GO website (http://geneontology.org/) using the PANTHER Overrepresentation Test (Mi et al. 2019). The TAIR10 identifiers of the transcripts in the ortholog-transcriptome were provided as reference list. The TAIR10 identifiers of the transcripts which were identified as significantly differently expressed genes by both programs, DESeq2 and EdgeR, were provided as analyzed list. *Arabidopsis thaliana* was chosen as organism and “GO Molecular function complete” was selected as annotation data set. Enriched GO categories were determined using the Fisher’s Exact Test with False Discovery Rate correction.

GO categories and terms were also determined using the AnnotationDbi in R. Transcripts associated with the term “DNA-binding transcription factor activity” were analyzed further.

### Promoter analyses

Binding of transcription factors to the promotors of genes involved in fruit opening was analysed using ChIP-Hub (http://www.chip-hub.org/). ChIP-Hub provides access to data on binding sites determined using chromatin immunoprecipitation followed by sequencing (ChIP-seq). On the ChIP-Hub website, *A. thaliana* was chosen as species and binding data was visualized on the WashU EpiGenome Browser. For each fruit development gene, 1,500 nucleotides upstream of the translation start codon were investigated and each occurance of binding of one of the transcription factors found to be differentially regulated was noted.

## List of abbreviations

AGL104: AGAMOUS-LIKE 104
ALC: ALCATRAZ
AP2: APETALA2
ARF: auxin-response factor
AS1: ASYMMETRIC LEAVES 1
AS2: ASYMMETRIC LEAVES 2
bHLH: basic helix-loop-helix
BP: BREVIPEDICELLUS
BRC1: BRANCHED1
BUSCO: Benchmarking of Universal Single-Copy Orthologs
ChIP-seq: chromatin immunoprecipitation followed by sequencing
DDEG: differently differentially expressed gene
DEG: differentially expressed gene
DZ: dehiscence zone
FIL: FILAMENTOUS FLOWER
FLC: FLOWERING LOCUS C
FUL: FRUITFULL
GO: gene ontology
IND: INDEHISCENT
JAG: JAGGED
LRT: Likelihood Ratio Test
MSG2: MASSUGU 2
MYB57: MYB DOMAIN PROTEIN 57
NF-YB2: NUCLEAR FACTOR Y-B2
NF-YB10: NUCLEAR FACTOR Y-B10
nt: nucleotide sequence database of NCBI
NTT: NO TRANSMITTING TRACT
NZZ: NOZZLE
OBP4: OBF BINDING PROTEIN 4
PHB: PHABULOSA
PIF1: PHY-INTERACTING FACTOR 1
RPL: REPLUMLESS
RVE6: REVEILLE 6
SHP1: SHATTERPROOF1
SHP2: SHATTERPROOF2
SPL: SPOROCYTELESS
SPL4: SQUAMOSA PROMOTER BINDING PROTEIN-LIKE 4
SPT: SPATULA
TCP4: TCP FAMILY TRANSCRIPTION FACTOR 4
TF: transcription factor
TRY: TRIPTYCHON
WOX13: WUSCHEL-RELATED HOMEOBOX gene 13
YAB3: YABBY3
ORE1: ORESARA1
ZFP2: ZINC FINGER PROTEIN 2

## Declarations

### Ethics approval and consent to participate

Not applicable

### Consent for publication

Not applicable

### Availability of data and materials

The datasets supporting the conclusions of this article are available at NCBI under the BioProject identifier PRJNA769250 https://www.ncbi.nlm.nih.gov/bioproject/769250.

### Authors’ contributions

GT conceived the project. LG and GT designed the experiments. KK performed the experiments. LG, NFP and MH analyzed the transcriptome data. LG, GT, MM and SR wrote the manuscript. All authors read and approved the final manuscript.

## Acknowledgements

We thank Heidi Küster for skillful technical assistance as well as Tobias Horst Rogalla and Georgia Daraki for preliminary analyses.

